# Protocol for using an ELISA to detect total α-synuclein levels in *Drosophila melanogaster* lines expressing human α-synuclein point mutations

**DOI:** 10.64898/2026.01.07.698009

**Authors:** Marco Sciortino, Raquel Velazquez, Haven Tillmon, Swati Banerjee

## Abstract

In Parkinson’s disease (PD), aggregation of alpha-synuclein (α-syn) contributes to neuronal dysfunction and death, particularly in dopaminergic neurons, driving disease progression. Several pathogenic point mutations have been identified in human α-syn such as *A30P, E46K, H50Q, G51D, A53T* and *A53E* that have been associated with PD. In this study, a sandwich ELISA assay was developed to quantify total α-syn levels in various *Drosophila melanogaster* genotypes expressing *A30P, E46K, G51D and A53T*. Using this approach, we compared α-syn levels across wild-type (WT) and mutant forms of the protein. The *E46K* and *A53T* mutations exhibited higher total α-syn concentrations compared to WT and the *G51D* mutation. In addition to characterizing mutationdependent differences in α-syn levels, this assay was applied to evaluate smallmolecule modulators that inhibit α-syn aggregation. These findings demonstrate that the ELISA-based approach provides a useful platform for quantifying α-syn and assessing potential therapeutic compounds targeting αsyn pathology.

## Before you begin

α-synuclein is a neuronal protein that aggregates under pathological conditions to form inclusions called Lewy bodies ^1,2^. These aggregates define a group of neurodegenerative disorders, including PD, collectively known as synucleinopathies. In PD, accumulation of α-synuclein contributes to motor and non-motor symptoms such as resting tremor, gait disturbances, cognitive decline, depression, and reduced quality of life ^3,4,5^.

Several pathogenic point mutations have been identified in human α-syn such as *A30P, E46K, H50Q, G51D, A53E and A53T*.^6,7,8,9^ In this study, we compare flies expressing three pathogenic α-syn variants (*E46K, G51D*, and *A53T*) with flies expressing wild-type α-syn. Using the ELISA assay kit, we quantified total human αsyn levels across each genotype. This ELISA assay kit was previously used in an *in vivo* drug-screening experiment in *Drosophila* designed to identify inhibitors of α-syn aggregation (*Sciortino and Banerjee, unpublished data*) and reliably detected changes in α-syn levels between untreated controls and drug-treated groups.

### Innovation

Existing methods for measuring α-syn levels, such as western blotting and immunohistochemistry, rely on relative signal intensity, lack precise quantification and require normalization to loading controls. Subtle yet biologically meaningful changes may be missed as signal intensity does not always scale linearly with protein concentration. In addition, differences in antibody quality, exposure times, staining procedures and imaging parameters across experiments or laboratories can introduce inconsistency thereby limiting reproducibility and statistical rigor.

This protocol introduces a sensitive ELISA-based assay that quantifies total human αsyn using a capture antibody and HRP-conjugated detection antibody. While the commercial ELISA kit is validated for mammalian brain homogenates and cell or tissue lysates prepared in RIPA lysis buffer, this protocol adapts the assay for *Drosophila melanogaster* fly head lysates prepared in 1% NP-40 lysis buffer. Due to differences in detergent composition and extraction conditions between RIPA and NP-40 buffers, optimization of homogenization volume, dilution factors, and sample preparation steps were required. This modification extends the applicability of the assay to smaller volume samples in addition to analyzing invertebrate models.

The assay also enables detection in the picogram range, reduces sample requirements, and improves quantitative accuracy, sensitivity, and reproducibility compared to traditional approaches. Additionally, the plate-based format supports scalable, high-throughput analysis, providing a key methodological advantage for quantitative α-syn measurement.

### Generating Drosophila Stock

🕓 **Timing:** a minimum of 2 weeks

1. Obtain *Drosophila* stocks from the Bloomington Drosophila Stock Center, Indiana University, Bloomington: https://bdsc.indiana.edu/index.html **Note:** A fly stock containing an *elav-GAL4* driver on the X chromosome (stock 458), in which the neuronal *elav* promoter drives broad GAL4 expression in postmitotic neurons. **Note:** A fly stock containing the *UAS-hSNCA* construct on the third chromosome (stock 8146), which expresses wild-type human α-syn (SNCA) under UAS control. **Note:** A fly stock containing the *UAS-hSNCA*.*E46K* construct on the second chromosome (stock 80043), which expresses human SNCA with familial Parkinson’s disease amino acid change, E46K, under UAS control. **Note:** A fly stock containing the *UAS-hSNCA*.*A53T* construct on the second chromosome (stock 95241) which expresses human SNCA with familial Parkinson’s disease amino acid change, A53T, under UAS control. **Note:** A fly stock containing the *UAS-hSNCA*.*G51D* construct on the second chromosome (stock 80044), which expresses human SNCA with familial Parkinson’s disease amino acid change, G51D, under UAS control. **Note:** The *UAS-hSNCA*.*A53T* (stock 95241) and *UAS-hSNCA*.*G51D* (stock 80044) stocks may contain flies carrying the *CyO* balancer chromosome, which produces a curly-wing phenotype. Balancer chromosomes are specially engineered chromosomes with multiple nested inversions that suppress recombination during meiosis and carry dominant visible markers such as curly wings. Most balancer chromosomes are lethal when homozygous.
  a. Maintain stocks in vials with 10 ml of standard fly food.
2. Set individual crosses with the *elav-Gal4* virgin females and males from the UAS fly lines described above.
  a. Place 20-25 CO_2_-anesthetized virgin females from the *elav-Gal4* stock in standard food vials with 10 anesthetized males from the UAS stock. **Note:** Collecting female flies within 6-8 hours of eclosion ensures that they have not been fertilized. For assistance in distinguishing males from females, we recommend *Atlas of Drosophila Morphology: Wild-type and Classical Mutants*^10^.
  b. Transfer the parent crosses to fresh food vials daily for 5 days.

i. First, gently tap the flies on a mouse pad to collect the flies at the bottom of the vial.
ii. Quickly remove the flug and invert the original vial over a new food vial.
iii. Seal the new vial with a fresh flug.

a. Keep the original vials with eggs, larvae and pupa from the F1 generation. **Note**: Adult flies typically begin to eclose from their pupal cases approximately 12 days after mating. The parent crosses should be discarded within 10 days to prevent inter-generation contamination.
b. Begin collecting flies from the F1 progeny from the day of eclosion (day 0) in new food vials. Collect both males and female flies of the desired genotypes (Figure 1). **Note**: If any of the UAS parent lines carry the *CyO* balancer chromosome, collect only the non-curly-winged progeny. These flies will have the correct genotype, carrying both the UAS transgene and the GAL4 driver.

**Figure 1.**
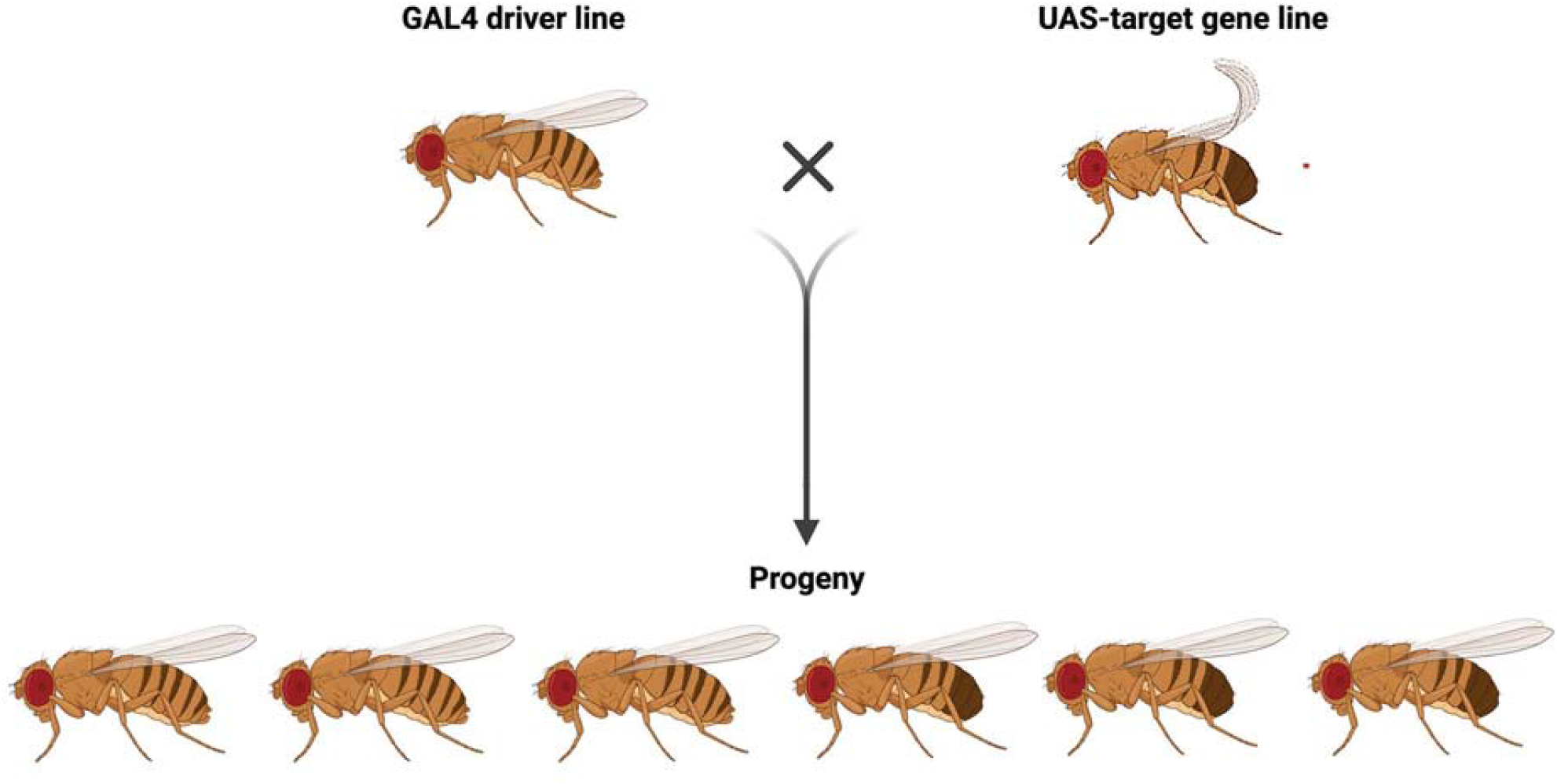
A crossing scheme of *Drosophila melanogaster* lines to produce F1 progeny. A cross between the GAL4 driver line with the UAS transgenic line that results into the F1 progeny carrying both the GAL4 driver and the UAS construct.

### Key resources table

| REAGENT or RESOURCE | SOURCE | IDENTIFIER |
| --- | --- | --- |
| **Chemicals, peptides, and recombinant proteins** |  |  |
| NP-40 | Fisher Scientific | 9016-45-9 |
| NaCl | Fisher Scientific | 7647-14-5 |
| HEPES | Millipore Sigma | 7365-45-9 |
| MgCl<sub>2</sub> | Millipore Sigma | 7791-18-6 |
| CaCl<sub>2</sub> | Fisher Scientific | 10035-04-8 |
| HPLC H<sub>2</sub>O | Fisher Scientific | 7732-18-5 |
| Roche cOmplete™, Mini, EDTA-free Protease Inhibitor Cocktail | Millipore Sigma | 11836170001 |
| **Critical commercial assays** |  |  |
| Sensolyte Anti- α-Synuclein (Human) ELISA kit | Anaspec | AS-55550-H |
| **Experimental models: Organisms/strains** |  |  |
| P{w[+mW.hs]=GawB}elav[C155] | Bloomington Stocks | 458 |
| w[*]; P{w[+mC]=UAS-Hsap\SNCA.F}5B | Bloomington Stocks | 8146 |
| w[*]; P{w[+mC]=UAS-hSNCA.E46K}2 | Bloomington Stocks | 80043 |
| w[*]; P{w[+mC]=UAS-SNCA.A53T.U}2 | Bloomington Stocks | 95241 |
| w[*]; P {w[+mC]=UAS-hSNCA.G51D}2 | Bloomington Stocks | 80044 |
| w[1118] | Bloomington Stocks | 5905 |
| **Software and algorithms** |  |  |
| Prism v.10 | GraphPad | graphpad.com |
| Microsoft Excel (version 16.89.1) | Microsoft Corporation | www.microsoft.com |
| SoftMax v. 7.1.2 | SoftMax | Moleculardevices.com |
| **Other** |  |  |
| Narrow Plastic Fly Food Vials | Archon Scientific | D20101- CASE |
| Flugs for Narrow Plastic Vials | Genesee Scientific | 49-102 |
| Square-bottom Fly Food Bottles | Archon Scientific | D20601- CASE |
| Flugs for Stock Bottles | Genesee Scientific | 49-100 |
| CO<sub>2</sub> Anesthetizing Pad | Genesee Scientific | 59-114 |
| Stemi 355 microscope | Zeiss | 435066-9530-000 |
| Flypad Frame | Genesee Scientific | 59-118 |
| Flystuff Blowgun | Genesee Scientific | 54-104 |
| FlyStuff Foot Pedal Complete System with Standard Flypad | Genesee Scientific | 59-121C |
| Flystuff Drosophila Trays and Dividers | Genesee Scientific | 32-122 |
| Gas Regulator | Agilent | 5183-4644 |
| Carbon Dioxide Cylinder | Linde Gas and Equipment Company | CD50 |
| Scalpel | Fine Science Tools | 10003-12 |
| Blade | Fine Science Tools | 10011-00 |
| Metal Probe | Fine Science Tools | 10140-01 |
| Micro Tissue Grinder (0.2 mL) | DWK Life Sciences | 885470-0000 |
| 1.7 mL microtubes | Axygen | MCT-175-C |
| Eppendorf Research Plus 8-channel Multichannel Pipette 120-1200 µL | Pipette.com | ES-8-1200-R |
| Pipetman L Starter Kit | Gilson | F167370 |
| Reagent Reservoir | Pipette.com | P8050 |
| Insulated Centrifuge | Fisher Scientific | p-8471006 |
| Ice Bucket Round 4L Blue 1CS | Government Scientific Source | 07210123 |
| Kimwipes | Fisher Scientific | 06-666 |
| SpectraMax iD3 | Molecular Devices | 5054747 K |

### Materials and equipment

#### 1% NP-40 extraction buffer composition

Materials and equipment 1% NP-40 extraction buffer composition
| Reagent | Final Concentration | Amount |
| --- | --- | --- |
| NP-40 | 1% | 100 µL |
| NaCl | 100 mM | 200 µL |
| HEPES (pH 7.5) | 50 mM | 500 µL |
| MgCl<sub>2</sub> | 1 mM | 10 µL |
| CaCl<sub>2</sub> | 1 mM | 10 µL |
| HPLC H<sub>2</sub>O | n/a | 9.18 mL |
| Protease Inhibitor | n/a | 1 tablet |
| **Total** |  | **10 mL** |

Protease inhibitor should be stored at 2°C to 8°C. All other reagents can be stored at 22°C to 25°C for at least one year.

### Step-by-step method details

#### Aging flies

The purpose of this step is to collect and age the flies.

🕓 **Timing:** 1 to 21 days

1. Collect an equal number of male and female flies resulting from the fly crosses described in step 2d (Generating *Drosophila* Stock).
2. Maintain flies in food vials, transferring them twice a week into fresh food vials until the flies reach 21 days of age.

### Homogenization preparation

This step ensures the homogenizers are ready for use before samples are introduced.

🕓 **Timing:** 10 min

3. Use a separate pestle and glass homogenization tube for each sample in the ELISA assay. **Note:** When selecting the pestle and tube, ensure the pestle makes direct contact with the tube to properly crush the fly heads.
4. Label each homogenizer with their respective genotype.
5. Pipet 40 microliters (µL) of NP-40 extraction buffer into each tube and place the tube and pestle on ice. **Note:** When placing the tube on ice, ensure that ice does not enter the tube.

### Collecting fly heads

The purpose of this step is to isolate fly heads of desired genotypes.

🕓 **Timing: ~**45 min

6. Take out the ELISA kit from the 4°C refrigerator and allow it to reach 23°C before use.
7. Anesthetize 3 male and 3 female flies of desired genotype per sample (n) on the CO_2_ fly pad.
8. Use a scalpel to cut the fly heads from the respective genotypes. **Note:** While cutting the fly heads, do not damage the fly eyes.
9. Transfer the fly heads into the chilled NP-40 extraction buffer using a probe. **Note:** When placing the heads into their respective tubes, ensure they are immersed in the NP-40 extraction buffer.
10. Repeat steps 7-9 for collecting samples from each genotype.

### Sample preparation

The purpose of this step is to prepare cell lysates for downstream analyses.

🕓 **Timing:** 1 hr

11. Using a Kimwipe, remove any condensation on the pestle. Crush the fly heads to ensure effective cell and tissue disruption during homogenization. **Note:** When crushing fly heads, make sure every head is homogenized to avoid inconsistencies between samples.
12. Turn on refrigerated centrifuge and set to 4°C
13. Let samples sit for 10 minutes on ice.
  a. Transfer 40 µL of each sample into a clean Eppendorf tube.
  b. Place samples in the centrifuge set at 4°C.
  c. Centrifuge at 18,400 x g for 15 minutes.
14. After centrifugation, pipet 30 µL of sample, without disturbing the pellet, into new Eppendorf tubes labeled with the respective genotypes and keep on ice. **Note:** If the pellet is disturbed, it is best to repeat the centrifugation step for 34 minutes.

### ELISA assay day one

The purpose of this step is to prepare the necessary components and incubate the samples and standards with the detection antibodies provided.

🕓 **Timing:** 1.5 hrs

**Note:** Prepare additional volume of all buffers/solutions to account for extra wells and pipetting loss.

15. Pipet 25 µL of sample into an Eppendorf tube labeled with the respective point mutation
  a. Add 75 µL of NP-40 extraction buffer to prepare a master dilution stock for each sample.
16. Mix each master dilution stock by inverting, and spin down before use.
17. Combine 498.75 µL of component C (wash buffer provided in the kit) along with 1.25 µL of the master dilution stock of each sample in a labeled Eppendorf tube to prepare the sample solution for each point mutation (Table 1).
18. Invert and spin down each sample to ensure proper distribution.
19. Suspend Component B in 1 milliliter (mL) of Component C at a concentration of 100,000 picograms (pg) per mL (Table 1).
  a. Invert tube gently several times, and let it sit at 23°C for 10-15 minute.
  **Note:** After adding 1 mL of Component C, use Component B to prepare the standards outlined in step 20 within one hour, and then apply to ELISA plate outlined in step 22.
20. Using serial dilution, prepare 9 standards for the ELISA assay. Standard 1 (10,000 pg/mL), standard 2 (500 pg/mL), standard 3 (250 pg/mL), standard 4 (125 pg/mL), standard 5 (62.5 pg/mL), standard 6 (31.25 pg/mL), standard 7 (15.625 pg/mL) and standard 8 (7.8125 pg/mL). **Note:** Only standards 2-8 should be used for the ELISA assay, as the protocol is optimized for the sample concentrations within this range. A successful standard curve should yield an R^2^ value ≥ 0.95. If this criterion is not met, repeat the assay using freshly prepared, high-quality standards.
21. Arrange and label the strips of wells (Component A) of the ELISA plate (Table 1).
  a. Reserve 14 wells for the standards, 3-5 wells per sample (n), and 3 wells for the blank readings (See Figure 2).
  **Note:** Although only 300 µL is required to load three replicate wells (100 µL per well), 500 µL of each sample is prepared in step 17 to allow for additional technical replicates or to repeat loading, if needed.
22. Add 100 µL of each standard or sample to the designated wells. Load standards in duplicate and samples in triplicate (See Figure 2).
23. Spin down component G, (the HRP-conjugated detection antibody) to ensure all contents can be retrieved from the bottom of the tube (Table 1).
  a. Dilute Component G 200-fold with component C.
  b. Add 50 µL of the resulting preparation to each well.
24. Cover the ELISA plate with the provided adhesive film and incubate at 4°C for 16 hours. **Note:** This allows α-syn in the samples to bind to the plate-coated capture antibodies and to simultaneously associate with the HRP-conjugated detection antibody.

**Table 1.** Components of Anaspec ELISA kit.

|  |  |
| --- | --- |
| Component A | Anti-α-synuclein 8-well strips |
| Component B | α-synuclein Standard, human (1 $\mu\text{g}$) |
| Component C | 1X Sample Dilution Buffer |
| Component D | 10X Wash Buffer |
| Component E | TMB Color Substrate Solution |
| Component F | Stop Solution |
| Component G | Detection Antibody |
| Component H | Adhesive plate covers |

**Figure 2.**
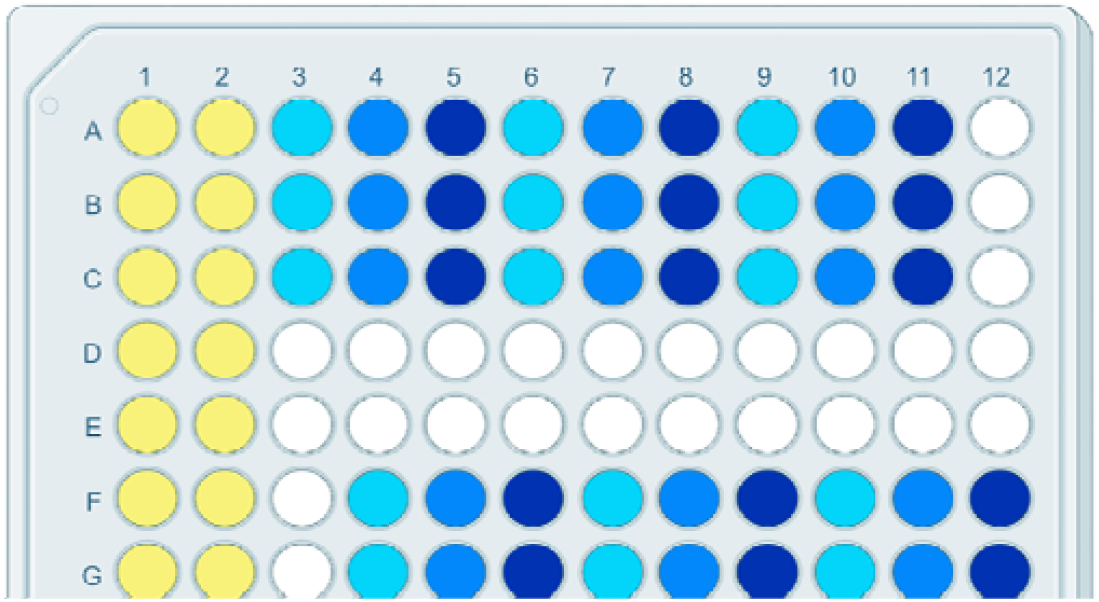
Representative ELISA plate layout. The figure shows the ELISA plate with the placement of standards (yellow), blank controls (grey), and experimental samples loaded in triplicate for each genotype across the 96-well plate. Figure created using BioRender.com.

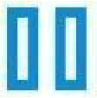 **Pause Point:** 16 hours

### ELISA assay day two

The purpose of this step is to perform washes and add the substrate solution before terminating the reaction.

25. Remove the ELISA assay kit components and allow them to reach 23°C.
26. Prepare the required amount (350 µL per well for six washes) of 1x wash buffer by diluting Component D (10X wash buffer) 1:10 with deionized (DI) water (Table 1).
27. Remove and discard adhesive film on the ELISA plate.
28. Remove all liquid from the ELISA plate by gently blotting it on paper towels.
  a. Wash each well six times with 350 µL of wash buffer.
  **Note:** Allow each wash to sit for 15 seconds before removing it.
29. Add 100 µL of Component E into each well.
  a. Incubate at 20°C for 18 minutes until a blue gradient is observed (Table 1).
  **Critical:** Cover the ELISA plate during incubation, as Component E is light sensitive.
30. Add 50 µL of Component F (stop solution) into each well. A color change from blue to yellow should be observed (Table 1). **Critical:** The plate must be read within 20 minutes of adding Component F to all the wells.

The expected outcome of this protocol is the quantification of total human α-syn concentrations (pg/mL) in *Drosophila* samples using an ELISA assay (Figure 3). Flies expressing human α-syn WT serve as a positive control, confirming that the assay reliably detects the α-syn protein. The *w*^*1118*^ genotype serves as a negative control, as these flies and *Drosophila* in general, do not express α-syn endogenously. Among the α-syn variants, flies expressing the *E46K* and *A53T* mutations are expected to exhibit higher α-syn concentrations compared to α-syn *WT* and the *G51D* mutation. In contrast, the *G51D* mutation is expected to show comparatively lower α-syn levels, suggesting reduced accumulation or stability of this variant. Together, these findings demonstrate that the ELISA assay can detect differences in α-syn levels in *Drosophila* carrying various disease-associated mutations in synucleinopathies.

**Figure 3.**
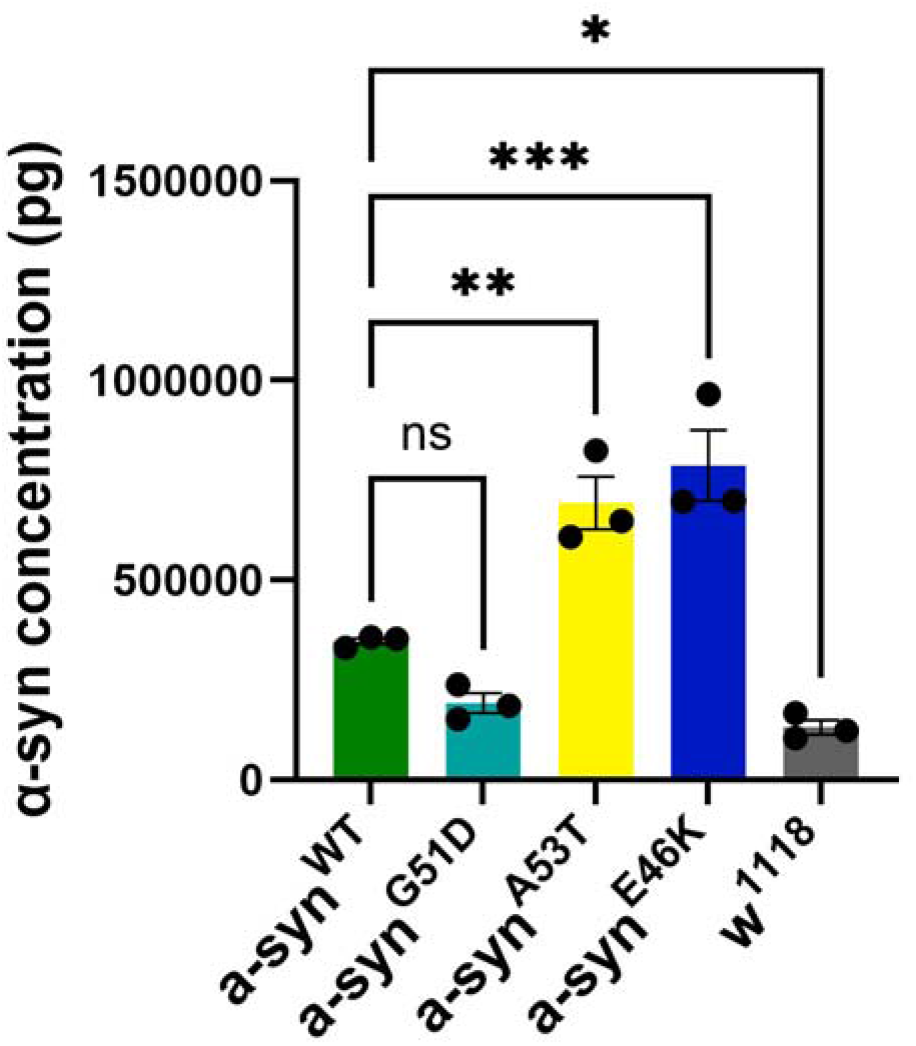
Quantification of human α-syn concentrations in specified genotypes of *Drosophila* by ELISA. Total α-syn levels (pg/mL) were measured from the brains of *Drosophila* expressing human α-syn WT and human α-syn point mutations, G51D, A53T, and E46K pan-neuronally using the *elav-Gal4* driver. The *w*^*1118*^ genotype served as a negative control. Data are presented as mean ± SEM from n = 3 biological replicates per genotype. Statistical comparisons were performed using one-way ANOVA followed by Dunnett’s multiple comparisons test. ns = not significant, *α-syn*^*WT*^ versus *α-syn*^*G51D*^; \*\**p=0*.*0027, α-syn*^*WT*^ versus *α-syn*^*A53T*^; \*\*\**p=0*.*0004, α-syn*^*WT*^ versus *α-syn*^*E46K*^; \**p=0*.*0459, α-syn*^*WT*^ versus *w1118*.

### Quantification and statistical analysis

31. Set the plate reader to measure absorbance at 450 nm, and include a 10s orbital shake before the reading.
32. Export the data into an Excel spreadsheet.
  a. Calculate the average of the standard replicates.
  b. Subtract the average of the three blank readings from each averaged standard.
33. Plot the averaged standard readings (y-axis) against their corresponding standard concentrations (x-axis) to generate the standard curve used to calculate α-syn concentrations in the samples (Figure 4).
34. Average the sample readings for each n.
  a. Subtract the average blank value to correct for background signal.
35. Calculate the sample concentration (pg/mL) using the slope and intercept from the standard curve as follows:

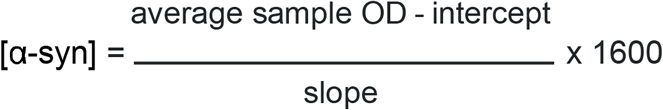 **Note:** The total dilution factor of 1600 accounts for the 1:4 NP-40 dilution (step 15a), and the subsequent 1:400 dilution (step 17).

**Figure 4.**
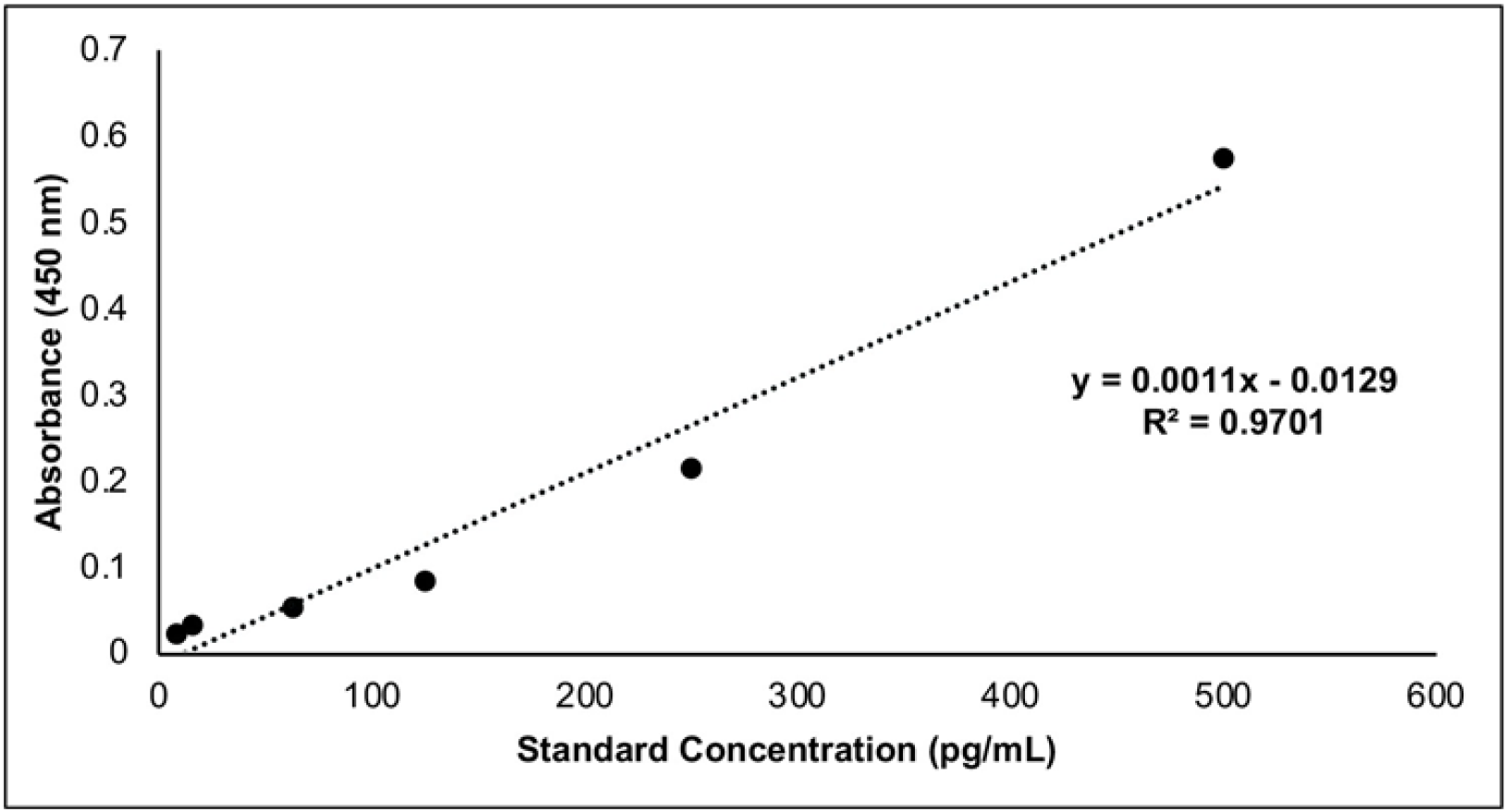
ELISA standard curve used to determine α-syn concentrations in the *Drosophila* samples. Known concentrations of α-syn standards (pg/mL) were plotted against absorbance values measured at 450 nm. A linear regression analysis was used to generate an equation for sample concentration calculation (R^2^ = 0.97).

### Limitations

This ELISA assay protocol is specialized for detecting total human α-syn levels and cannot be used to detect aggregated forms of α-syn that may exist in pathological conditions such as oligomeric and fibrillar forms. In addition, this protocol is optimized for *Drosophila melanogaster* and has not been tested with other animal models.

### Troubleshooting

#### Problem 1

Low signal detection for standards and samples (related to steps 15-18, 22 and 23a).

#### Potential solution

Verify that the 16-hour incubation at 4°C is performed correctly, as incorrect incubation time or temperature reduces binding between α-syn, the capture antibody, and the HRP-conjugated detection antibody. Ensure that the detection antibody was diluted correctly according to step 23a (ELISA assay day one). If the signal intensity remains low, increase the amount of starting material to raise the total protein concentration.

#### Problem 2

High background signal affecting data sets (related to step 28a).

#### Potential solution

Ensure that all wells are washed 6 times with 1x wash buffer and blotted on paper towels after each wash.

#### Problem 3

The standard curve R^2^ value is too low (related to step 19, 20 and 22).

#### Potential solution

Properly vortex and spin down each standard before and after every serial dilution, as incomplete mixing can result in a non-linear standard curve. Ensure Component B is fully dissolved and equilibrated to 23°C for 10-15 minutes, as incomplete dissolution can lead to inaccuracies during serial dilution.

## Resource availability

### Lead contact

Further questions and information for reagents and resources should be directed and will be fulfilled by the lead contact, Swati Banerjee.

### Technical contact

Technical issues can be directed to the technical contact, Marco Sciortino.

### Materials availability

This study did not generate new reagents.

### Data and code availability

This study did not analyze/generate datasets/code.

## Acknowledgments

We thank the Bloomington *Drosophila* Stock Center, Indiana University, for the *Drosophila* stocks used in this study. This work was supported by funds from the National Institutes of Health/National Institute of Neurological Disorders and Stroke grant R01NS134867, the Perry and Ruby Stevens Parkinson’s Disease Center of Excellence and the Long School of Medicine, UT San Antonio Health Science Center.

## Author contributions

M.S. developed the protocol, conducted the experiments, analyzed the data and wrote the original draft of the manuscript. R.V. conducted experiments to validate the findings. H.T. helped with optimizing the protocol and data analysis. S.B. conceptualized and supervised the project, provided resources and acquired funding, reviewed and edited the manuscript, and communicated with the journal. All authors contributed to manuscript review and editing.

